# Diminished motor neuron activity driven by abnormal astrocytic GLAST glutamate transporter activity in spinal muscular atrophy is not fully restored after lentiviral SMN delivery

**DOI:** 10.1101/2022.05.30.494049

**Authors:** Emily Welby, Allison D. Ebert

**Affiliations:** Department of Cell Biology, Neurobiology and Anatomy, Medical College of Wisconsin, Milwaukee, WI, USA

**Keywords:** survival motor neuron (SMN), human induced pluripotent stem cells, EAAT1/GLAST, EAAT2/GLT1, synapse, caveolin-1, microelectrode array, lentiviral gene therapy

## Abstract

Spinal muscular atrophy (SMA) is a pediatric neuromuscular disease characterized by the loss of the lower spinal motor neurons due to survival motor neuron (SMN) deficiency. Motor neuron dysfunction at the glutamatergic afferent synapse is observed during early stages of SMA disease progression, which could be targeted therapeutically prior to cell death. However, the motor neuron cell autonomous and non-cell autonomous disease mechanisms driving this phenotype remain unclear. Our study reveals a non-cell autonomous SMN-associated disease mechanism affecting glutamate transporter (GLAST) activity in astrocytes that contributes to human motor neuron dysfunction in SMA. Transcriptomic analysis of SMA patient human induced pluripotent stem cell (iPSC)-derived astrocytes identified a significant downregulation of genes associated with astrocytic regulation of the synapse, including glutamate neurotransmission. This finding was substantiated by our microelectrode array analysis of motor neuron activity, which was severely diminished specifically in the presence of patient-derived astrocytes. Co-culturing patient-derived motor neurons with healthy-derived astrocytes showed comparable firing rates and bursting activity to healthy-derived motor neurons, suggesting diminished neural activity is an astrocyte-mediated phenotype in this system. Towards defining astrocyte-intrinsic defects that could induce motor neuron dysfunction, we identified abnormally low levels of excitatory amino acid transporter (EAAT1/GLAST) in patient-derived astrocytes, which when selectively inhibited in healthy co-cultures could phenocopy the diminished neural activity previously observed in patient-derived co-cultures. Caveolin-1, an SMN-interacting lipid raft protein associated with glutamate transporter regulation, showed increased protein levels and accumulation in patient astrocytes. Both GLAST and caveolin-1 phenotypes could be partially rescued via lentiviral-mediated SMN re-expression in patient astrocytes. Together, our work defines a novel SMN-associated disease mechanism involving abnormal glutamate transporter activity and regulation in astrocytes that can directly diminish motor neuron function in SMA.

## Introduction

Motor neuron synaptic dysfunction is one of the first pathological features observed during disease progression of the pediatric neuromuscular disease, spinal muscular atrophy (SMA). Specifically, lower alpha spinal motor neurons responsible for proximal and axial muscle innervation as part of the sensory-motor circuitry are particularly vulnerable to degeneration, leading to eventual muscle wasting, respiratory failure and premature death (1-3). Survival motor neuron (SMN) deficiency caused by the homozygous mutation of *SMN1* and lack of functional SMN protein generation from the *SMN1* paralogue, *SMN2* (4, 5), has been linked to morphological and functional synaptic defects affecting excitatory proprioceptive innervation of motor neurons within the spinal cord and peripheral neuromuscular junctions (NMJ) (6-14). Since motor neuron cell death and dysfunction are thought to be mechanistically uncoupled with synaptic defects preceding motor neuron cell death (15, 16), correction of neuronal activity during early stages of SMA disease progression in combination with currently available *SMN1/SMN2* treatments would be of high therapeutic value to decrease therapeutic variability observed in SMA patients (17).

Excitatory proprioceptive synaptopathy is thought to be one of the most consistent early phenotypes observed across various SMA animal models, prior to synaptic loss and NMJ dysfunction (9-11, 15, 18). Defects at the central excitatory synapses have been linked to SMA motor neuron hyperexcitability characterized by abnormally increased membrane excitability, affecting motor neuron action potential properties and firing rate frequencies (10, 16, 19-21). SMN re-expression specifically within SMA motor neurons can reverse the hyperexcitability phenotype and restore excitatory afferent synapses (11, 19, 20, 22); however, motor neuron SMN restoration only lead to modest improvements in survival rates and motor behavior *in vivo* (23). Other lines of evidence demonstrate that early motor neuron hyperexcitability and dysfunction is instigated by SMN deficiency in proprioceptive neurons (10) and interneurons (16) thought to be driven by decreased presynaptic glutamate (VGLUT1) neurotransmission (24). Specific SMN re-expression within sensory neurons not only improves excitatory neurotransmission at the afferent motor neuron synapses, but also improves NMJ functioning and motor behavior in several models of SMA disease (24, 25). This is highly suggestive that other CNS cell types may be contributing to motor neuron dysfunction. Astrocytes also play a fundamental role at these excitatory synapses regulating synapse formation and function (26-28) through the expression of plasma membrane proteins (e.g. ion channels, neurotransmitter receptors, cell adhesion molecules) or secretion of matricellular molecules (e.g. thrombospondins, gliotransmitters and microRNAs) into the synaptic cleft (29). It remains to be fully elucidated if astrocytes directly contribute to central excitatory synaptopathy in SMA.

Astrocyte reactivity and dysfunction are commonly observed across SMA patient-derived samples and SMA mouse models (15, 30, 31), including decreased neurotrophic factor secretion (32) and abnormally increased microRNA (miR-146a) production (33), which have a detrimental impact on neuronal health. In relation to motor neuron synaptic dysfunction, SMA mouse-derived tissues and SMN siRNA cell cultures models demonstrate reduced astrocytic potassium channel (K_ir_4.1) activity, excitatory amino acid transporter 1 (EAAT1/GLAST) levels (34) and ephrin expression (35). Since astrocytic SMN restoration could partially restore of VGLUT1+ bouton density at proprioceptive synapses and motor neuron neurotransmission at the NMJ (30), astrocyte intrinsic disease mechanisms likely correlate with SMN levels. We have begun to address astrocyte-mediated motor neuron dysfunction in human SMA disease by recently demonstrating decreased motor neuron activity and expression of synaptic-associated extracellular matrix genes (36) after miR-146a mimic treatment. Abnormal calcium signaling in patient iPSC-derived astrocytes (31) may also impact the ability of astrocytes to maintain neuronal synaptic support, but further work is needed to unravel these disease mechanisms in SMA.

While SMN has an established role in assembling small nuclear ribonucleoproteins (snRNP) and in pre-mRNA splicing, it remains to be fully established how SMN deficiency can lead to synaptic dysfunction in SMA. Reduced SMN levels may contribute to neuronal dysfunction via defects in neurite development, growth cone dynamics, cytoskeletal processes involved in axon elongation and RNP granule transportation (37-44). Recently, endosomal trafficking perturbation in SMN deficiency was shown to cause loss of synaptic vesicle docking in motor neurons, but also the disruption of endocytosis in non-neuronal cell types (45). Recent evidence demonstrates the direct interaction of SMN with ribosomal proteins and caveolin-1 within cholesterol-rich lipid rafts at the plasma membrane (caveolae) where it contributes to regulation of local protein translation during membrane remodeling and actin filament rearrangement (46, 47). Local protein translation of RNA binding protein (RBP)-bound mRNAs is a critical mechanism to meet synaptic demand and activity, including modulating neurotransmitter receptor and transporter turnover (48). Therefore, SMN acting as a molecular chaperone to facilitate proper RBP-mRNA-cytoskeleton interactions and transportation, local protein translation and caveolae-associated endocytosis may therefore have important implications for neuronal and astrocytic neurotransmission modulation when SMN protein levels are perturbed.

In this study, we provide further evidence that a non-cell autonomous disease mechanism involving glutamate transporter activity within SMA patient iPSC-derived astrocytes has a detrimental effect on motor neuron activity. Regardless of motor neuron iPSC derivation, severely diminished neural activity was consistently observed in co-cultures containing patient-derived astrocytes. As suggested by our transcriptomic data, patient astrocytes have a significant reduction in transcripts related to synaptic integrity, synaptic structure and ion buffering, but also glutamate neuromodulation. Specifically, we validated the severe reduction of excitatory amino acid transporter 1 (EAAT1/GLAST) protein levels in patient astrocytes. Interestingly, attempting to mimic reduced GLAST levels via specific inhibition of GLAST functioning in healthy-derived astrocyte co-cultures resulted in diminished motor neuron activity, comparable to the levels observed in patient-derived astrocyte co-cultures. We postulate that the abnormally upregulated levels of caveolin-1, a lipid raft scaffold protein with known associations with glial glutamate transporter regulation (49), may contribute to the reduced GLAST levels observed in patient-derived astrocytes. The partial rescue of GLAST and caveolin-1 phenotypes following lentiviral delivery of SMN to patient astrocytes suggests that this disease mechanism is associated with SMN deficiency.

## Material and Methods

### Human induced pluripotent stem cell lines and spinal cord tissue

Human induced pluripotent stem cell lines (iPSCs) from 2x SMA Type 1 patients (patient line 1 (7.12) and patient line 3 (8.2), 1x SMA Type 2 patient (patient line 2 (3.6), previously classified as Type 1) and 2x healthy individuals (healthy line 1 (21.8) and healthy line 2 (4.2); 4.2 is the parent of 3.6) were used in this study. All patient lines have 3 copies of SMN2. Frozen healthy and SMA lumbar spinal cord tissue was obtained from the University of Maryland Brain and Tissue Bank under the auspices of the NIH NeuroBioBank. The use of iPSCs and human spinal cord tissue was approved by the Medical College of Wisconsin Institutional Review Board (PRO00025822), the Institutional Biosafety Committee (IBC20120742) and the Human Stem Cell Research Oversight Committee.

### iPSC maintenance and differentiation

iPSC lines were cultured on Corning® Matrigel® Growth Factor Reduced Basement Membrane Matrix or Geltrex ™ LDEV-Free, hESC-Qualified, Reduced Growth Factor Basement Membrane Matrix (Gibco) coated 6-well plates in Essential 8 medium (Gibco) and were passaged every 3 days using Versene (EDTA-based, Gibco) when 80% confluent. iPSCs were differentiated into spinal cord-like motor neurons or astrocytes using protocols recently described in (50). Neural progenitor cells (NPCs) in the form of embryoid body aggregates (for motor neuron differentiation) or monolayer cultures (for astrocyte differentiation) were generated from iPSCs through dual SMAD inhibition (SB 431542 and LDN 1931899 (biogems) and Wnt activation (Chir 99021 (biogems). NPCs were subsequently pattered into a ventral-caudal cell fate using retinoic acid (100nM, Sigma) and smoothened agonist (500nM, biogems). Motor neuron aggregates were dissociated and plated down on to Matrigel® coated plates with maturation media supplemented with 10µM DAPT (biogems), 10ng/ml BDNF (Peprotech) and 20ug/ml GDNF (Peprotech) for terminal differentiation. NPC monolayers were passaged once a week using Accutase (STEMCELL Technologies), until passage 4 when NPCs were seeded at 15,000 cells/cm^^2^ on to Matrigel® or Geltrex ™ coating flasks in ScienCell Astrocyte medium containing 2% B27 to initiate astrocyte differentiation. Astrocyte cultures were passaged when confluent once a week and were considered fully differentiated by P4. Astrocytes used for mRNA sequencing were generated from EZ sphere cultures (51), which were maintained in poly-HEMA (Sigma) coated flasks T75 flasks in EFH100 medium (Stemline (Sigma), 100ng/ml EGF (Miltenyi Biotec Inc), 100ng/ml FGF (biogems), 5ug/ml heparin (Sigma) and passaged once a week using a tissue chopper. Passage 6-10 spheres were then dissociated, plated in medium containing 2% B27, and passaged weekly for 6 weeks prior to RNA collection.

### RNA extraction, mRNA sequencing and DeSeq2 analysis

Astrocyte cell pellets were processed for mRNA sequencing as previously described (36). Briefly, mRNA was extracted from astrocyte cell pellets using the Qiagen RNeasy kit with on-column DNaseI treatment. ERCC Spike-In Controls (Invitrogen) were added to 100ng of RNA per sample prior to processing with the NEBNext Poly(A) mRNA Magnetic Isolation Module (NEB #E7490) and NEBNext Ultra II RNA Library Prep Kit for Illumina (NEB # E7770) according to manufacturer’s protocol. Unique identifiers for each sample cDNA library were created using the NEBNext Multiple Oligos for Illumina Set kits. The DNA 100 Chip kit and Tapestation system (Agilent) was used to assess the quality of cDNA libraries generated. Single and pooled libraries were assessed using the NEBNext Library Quant kit for Illumina (NEB #E7630), Bio-Rad CFX384 real time thermocycler and calculated using the NEB qPCR webtool (https://nebiocalculator.neb.com/qPCR). Pool libraries were denatured and diluted following the Illumina NextSeq system Denature and Dilute Libraries Guide. Samples were sequenced with paired end reads on the Illumina NextSeq system. FASTQ files were processed using Expression count (STAR) pipeline in Basepair Tech before performing principal component analysis and DeSeq2 RNA seq analysis. The g:Profiler web server (52) was used to perform gene ontology analysis. All data have been made publicly available through NCBI Sequence Read Archive.

### Microelectrode array preparation, recordings and analysis

For co-culture analysis, motor neuron differentiation cultures (Day 14) and astrocyte cultures (P4+) were simultaneously dissociated and combined together in 1:1 motor neuron maturation medium and ScienCell Astrocyte Medium (with 2% B27) at a cell density ratio of 5:1 (24,000 motor neurons: 5,000 astrocytes per well). For motor neuron or astrocyte only wells, cells were seeding at the according densities and cultured in the same 1:1 co-culture media. Cells were plated on to poly-L-lysine (Sigma)/laminin (Sigma) coated 48 well CytoView microelectrode array (MEA) plates (Axion Biosystems) in a 10µL droplet directly over the recording electrodes of each well. Cells were left to attached for 1 hour at 37°C before carefully flooding the wells with 300µl of 1:1 co-culture media. Media was aspirated and replaced every other day. Spontaneous and electrically evoked motor neuron activity was recorded every other day for approximately 1 month (28 days) starting from Day 7 post-plate down, typically when spiking activity begun. MEA recordings were performed by the 1^st^ generation Maestro system (Axion Biosystems). Cells were rested for 10 minutes on the pre-heated (37°C) MEA stage prior to starting the 4-minute recordings. For the GLAST inhibitor (50µM UCPH101) treatment, recordings were capture before treatment and after a 15-minute incubation period with the inhibitor compound at 37°C. Post-treatment, cells were washed 3x with medium to promote drug wash out. Motor neuron activity was recorded across multiple days, as effects of inhibitor treatments were temporary. Version 2.4.2.13 of the AxIS acquisition software (Axion Biosystems) was used to record activity across 16 electrodes per well with a sampling frequency of 12.5 kHz and a digital high pass filter of 5Hz IIR. The spike-detecting threshold was defined as voltage exceeding 6 standard deviations away from the mean background noise. Butterworth digital filter settings with high (200 Hz) and low (3kHz) pass cuttoff frequencies were applied during recordings. 5 spikes/minute was the threshold used to determine an active electrode. For electrically evoked recordings, the Neural Stimulation with Artifact Elimination (0.5V for 400µs (36400µs total operation time) was applied to the cells by stimulating 1 electrode per well once every 10 seconds. Statistic Compiler files and Spike Detector files were extracted from raw AxIS (spontaneous recordings) and raw AxIS Artifact Eliminator files (electrically evoked recordings) for downstream analysis of weighted mean firing rate and bursting activity. Additional background noise determined by low levels of activity from astrocyte only wells were also removed from the electrically evoked analysis. The NeuralMetricTool (Axion Biosystems) was used to generate raster plots.

### Glutamate uptake assays

Astrocytes were seeded into Matrigel®-coated 96 well plates at a density of 10,000 cells per well and cultured overnight (37°C, 5% CO2). Treatment conditions included 100µM glutamic acid (Sigma), 100µM glutamic acid with 50µM of the selective GLAST inhibitor, UCPH101 (Abcam), or 100µM glutamic acid with 50µM/100µM of the selective GLT-1 inhibitor, dihydrokainic acid (DHK; Abcam) diluted in ScienCell Astrocyte Medium from compound stock solutions. Treated and untreated (astrocyte medium only) astrocytes were collected and processed after a 2-hour incubation period to determine intracellular glutamate levels using the Glutamate-Glo™ Assay (Promega) according to manufacturer’s instructions. This included lysing the cells with the Inactivation Solution (0.6N HCl; Fisher Scientific), before neutralizing the solution with 1M Tris Base (Sigma) and proceeding with the bioluminescent assay. Cells were thoroughly washed with PBS post-treatment and pre-lysis to remove any contaminating extracellular glutamate. Samples were run in technical duplicates on white Nunc™ 96 well assay plates (Thermo Scientific) with glutamate standards (0-50µM). Luminescence signal was detected and acquired using a GloMax ® luminometer (Promega). A standard curve was generated to determine glutamate concentrations of sample luminescence signal.

### Immunocytochemistry

Astrocytes were seeded at a density of 30,000 cells on to 1.5oz glass coverslips (Electron Microscopy Sciences), which were pre-coated with Matrigel® or Geltrex ™. For SMN and CAV-1 immunostainings, cells were fixed in ice cold 1:1 acetone/methanol for 5 minutes at 4°C. For GLAST immunostaining experiments, cells were fixed in 4% paraformaldehyde (PFA) for 12 minutes at room temperature. Coverslips were washed thrice with 1x PBS and stored in PBS at 4°C if immunostaining could not be performed immediately. Coverslips were processed within a few days of fixation. Cells were permeabilized in 0.2% Triton X-100 for 10 minutes, before blocking in 10% normal donkey serum for 1 hour. Cells were incubated overnight in primary antibody solution (0.1% Triton X-100 and 5% normal donkey serum) at 4°C. The following primary antibodies were used in this study: chicken anti-vimentin (1:400, Abcam), mouse anti-GLAST (1: 300, Miltenyi Biotec), mouse anti-caveolin1 (1:50, BD Biosciences), mouse anti-SMN (1:200, BD Biosciences), rabbit anti-FLAG (1:400, Sigma), rabbit anti-GFP (1:400, Cell Signaling). Cells were washed 3x with 1x PBS (5 minutes per wash) before incubating with secondary antibodies at room temperature for 1 hour. Secondary antibodies used in this study included donkey anti-mouse Alexa Fluor 488 (Invitrogen), anti-rabbit Alexa Fluor 568 (Invitrogen) and anti-chicken Alexa Fluor 647 (Invitrogen). Primary antibodies were omitted from secondary only controls. Cells were washed 3x with 1x PBS (5 minutes per wash) prior to 4’,6-diamidino-2-phenylindole (DAPI; 1:3000; Thermo Fisher) incubation for 5 minutes at room temperature to stain nuclei. Cells were washed 3x with 1x PBS (5 minutes per wash) and were mounted onto Superfrost Plus microscope slides (VWR) in VECTASHIELD antifade mounting medium (Vector Laboratories). Coverslips were sealed with nail polish and stored at 4°C prior to imaging.

### Microscopy

Z-projection images were acquired of astrocyte-stained coverslips using the Zeiss LSM980 confocal laser scanning microscope with AiryScan 2 and ZEN 3.4 (blue edition) acquisition software (Zeiss). Brightfield and epi-fluorescent images of motor neuron and astrocyte co-cultures on MEA plates were acquired using an inverted Nikon microscope and QCapture camera. Images were processed using ZEN 3.4 and ImageJ software.

### Western blot

Astrocytes cell pellets were lysed and sonicated in Pierce™ RIPA buffer (Thermo Scientific). Protein concentration (mg/ul) was then determined via Pierce™ BCA Protein Assay (Thermo Scientific) using BSA standards. 10µg of protein per sample was loaded into 4-15% Mini-PROTEAN® TGX Stain-Free™ Protein Gels (Bio-Rad) and run at 105V for 90 minutes. Protein samples then were transferred to PVDF membranes (Li-COR) at 105V for 60 minutes, which were dried overnight to facilitate protein crosslinking. After methanol rehydration, membranes were incubated in Revert™ 700 Protein Stain (Li-COR) and imaged using an Odyssey Scanner (Li-COR). After removal of the total protein stain using the Revert™ Destaining Solution (Li-COR), membranes were incubated with 1:1 dilution of Odyssey Blocking Buffer (Li-COR) and 1x TBS at room temperature for 1 hour.

Primary antibodies were diluted in the 1:1 blocking buffer with 0.2% Tween-20 and incubated overnight at 4°C. Primary antibodies for immunoblotting were used at the following dilution: mouse anti-caveolin1 (1:2000, BD Biosciences), rabbit anti-GFP (1:1000, Cell Signaling). Following 3x washes with 1x TBST, membranes were incubated for 30 minutes at room temperature in secondary antibody solution (1:1 blocking buffer with 0.2% Tween-20, 0.02% SDS, 1:8000 secondary antibody). Secondary antibodies used in this study included: IRDye 800 CW Donkey anti-mouse and IRDye 680 CW Donkey anti-rabbit (Li-COR). Blots were imaged using the Odyssey Scanner and protein band intensities were converted into grayscale and quantified using Image Studio (Li-COR) software. Antibody signal was normalized to the total protein REVERT stain according to manufacturer’s instructions.

### SMN lentivirus preparation and infection

Lentiviral DNA constructs containing FLAG tagged human *SMN1* (hEF1α:SMN-FLAG) or a GFP reporter downstream of human *SMN1* (pSN1-CMV:hSMN-hEF1α:GFP) were kindly provided by the laboratories of Dr. Xue Jun Li (University of Illinois-Chicago) and Dr. Christian Lorson (University of Missouri), respectively. Lentivirus generation was performed by the Viral Vector Core at the Medical College of Wisconsin/Blood Research Institute. Viral titers used in this study were 7.36E+08 TU/ml (hEF1α:SMN-FLAG) and 6.75+08 TU/ml (pSN1-CMV:hSMN-hEF1α:GFP). SMA patient-derived astrocytes were transduced at a MOI of 10 with 8ug/ml polybrene for 48 hours.

### Statistics, rigor and reproducibility

For each experiment, cultures were derived from 2 healthy individual iPSC lines and 3 SMA patient iPSC lines (N=5). For SMN re-expression experiments, 4 differentiation replicates derived from the 3 SMA patient iPSC lines were transduced with SMN:FLAG (2 reps per line) or SMN:GFP lentiviruses (2 reps per line;12 SMN rescue lines total). Data from glutamate transporter treatment experiments (DHK glutamate assay, MEA analysis, glutamate assay on SMN transduced astrocytes) were from 2 independent replicates with a minimum of 3 biological replicates in each experiment. At least 3 independent replicates with a minimum of 3 biological replicates in each were completed for all other experiments. Statistical significance within the mRNA sequencing data was determined as - log10 adjusted p-value < 0.05, log2 fold change <-2, >2. One-way or two-way ANOVA with Tukey’s multiple comparison test were performed to determine statistical significance (p-value<0.05) within glutamate assay and immunoblot experiments. Kruskal-Wallis test with Dunn’s multiple comparisons test was performed on MEA datasets to determine statistical significance (p-value<0.05). Statistical analysis was performed with GraphPad Prism software.

## Results

### Downregulation of genes involved in synapse regulation in SMA patient iPSC-derived astrocytes

To assess transcriptomic differences attributed to SMN deficiency in human astrocytes, mRNA sequencing was performed on healthy and SMA patient iPSC-derived astrocytes (Figure 1). Astrocytes used for these experiments were derived from EZ sphere cultures (51), which produced a forebrain-like astrocyte population. Principal component analysis revealed the separation of healthy and SMA datasets based on PC1 (72.50%), with some degree of variability noted between technical replicates (PC2: 16.43%; Figure 1A). A total of 2995 genes were statistically differentially expressed (adjusted p-value < 0.05) between healthy and patient-derived astrocytes (Figure 1B), with a higher number of downregulated genes observed in the patient-derived astrocytes (1734 genes, log2 fold change <-2) compared to the upregulated gene set (1221 genes, log2 fold change >2). Gene ontology analysis revealed a significant enrichment of biological process, cellular component and molecular function terms related to the synapse, synaptic signaling, and ion channel activity (Figure 1C) within the downregulated gene dataset. Many of these genes encode astrocytic-specific proteins with defined roles in synaptic ion buffering (e.g. K_ir_4.1 inwardly-rectifying channels, Na^+^ K^+^ ATPases; Figure 1D), regulating and maintaining structural synaptic integrity (e.g. cell adhesion molecules, ephrins and neurexin-neuroligin signaling; Figure 1D) and modulating glutamate-specific neurotransmission, including metabotropic and ionotropic glutamate receptor subunits, and excitatory amino acid transporters, EAAT1/GLAST (SLC1A3) and EAAT2/GLT-1 (SLC1A2) (red data points Figure 1B; Figure 1D).

**Figure 1.**
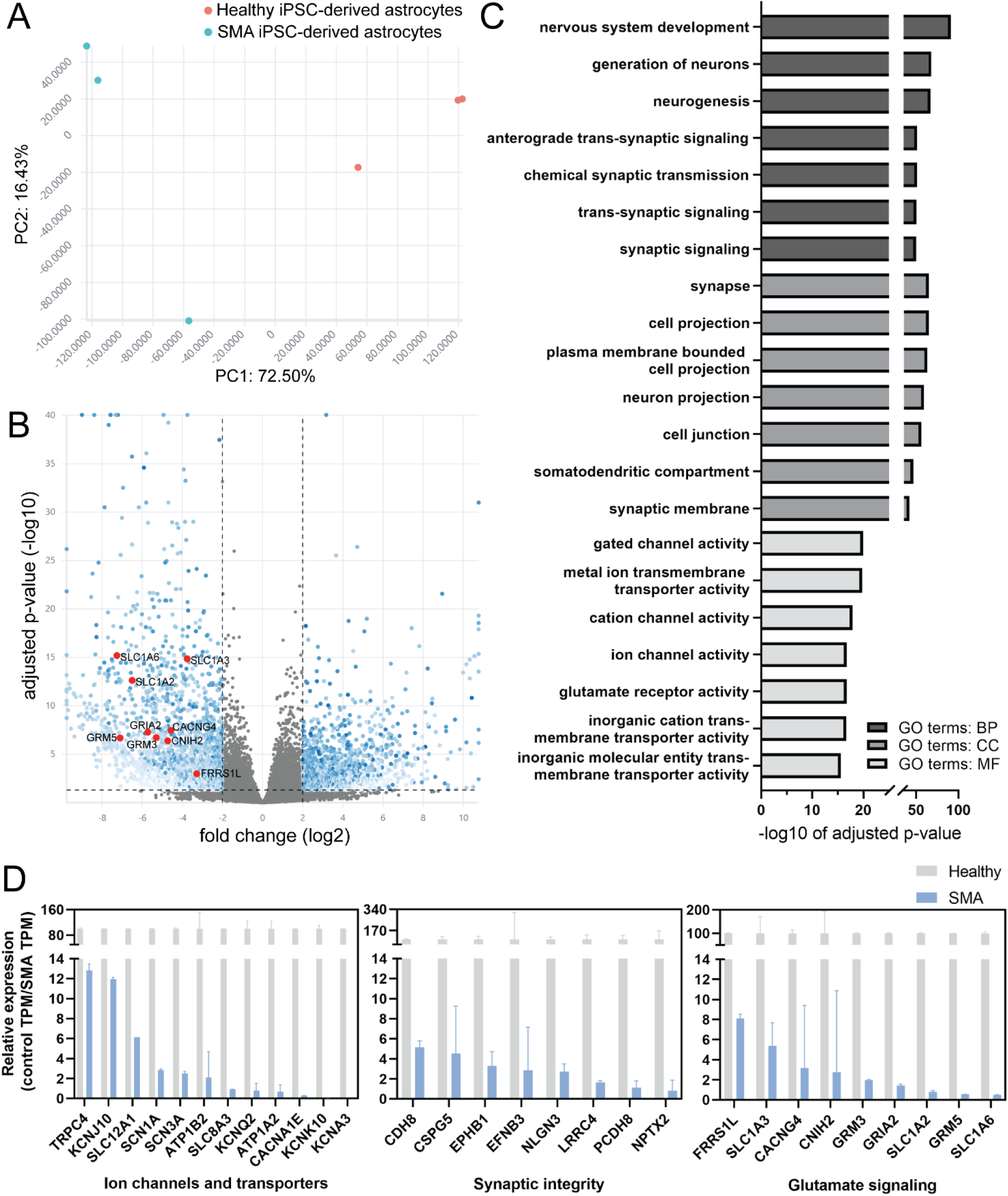
Significantly downregulated genes in SMA iPSC patient-derived samples are associated with astrocytic regulation of the synapse. **(A)** Principal component analysis of healthy (red datapoints) and SMA (blue datapoints) iPSC-derived astrocytes. Data are from 3 technical replicates per condition (n=3) from 3 independent cell lines (N=3). **(B)** Volcano plot showing significantly upregulated (-log10 adjusted p-value < 0.05, log2 fold change >2) and downregulated (- log10 adjusted p-value < 0.05, log2 fold change <-2) genes in SMA patient-derived astrocytes from the DeSeq2 differential gene expression analysis (healthy vs SMA samples). Red highlighted datapoints are downregulated genes related to astrocyte glutamate modulation. **(C)** Significantly (- log10 adjusted p-value) enriched biological process (BP), cellular component (CC) and molecular function (MF) terms from gene ontology analysis of genes downregulated in SMA astrocyte samples. **(D)** Relative expression values (transcripts per million; TPM) of downregulated genes in SMA astrocyte samples related to ion channels and transporters, synaptic integrity and glutamate signaling. For each gene, TPM values from SMA (blue bars) derived astrocytes were normalized to healthy (grey bars) TPM value to calculate relative expression, which is represented as a percentage and standard deviation.

### SMA patient iPSC-derived spinal cord astrocytes diminish motor neuron activity in direct contact co-cultures

We next wanted to determine if human spinal cord-like astrocytes, a more disease relevant cell population for studying SMA pathology, similarly display intrinsic defects in synaptic neuromodulation which could directly impact motor neuron activity. Ventral-caudal patterned motor neurons (53) and astrocytes (50) were derived from healthy and patient iPSCs before seeding together onto microelectrode array (MEA) plates (Figure 2A). The differentiation protocols used in this study allowed for the initial, independent generation of motor neurons and astrocytes from iPSC lines. Therefore, different motor neuron and astrocyte combinations could be co-cultured together to assess if changes in neural activity are motor neuron-intrinsic or driven by diseased astrocytes.

**Figure 2.**
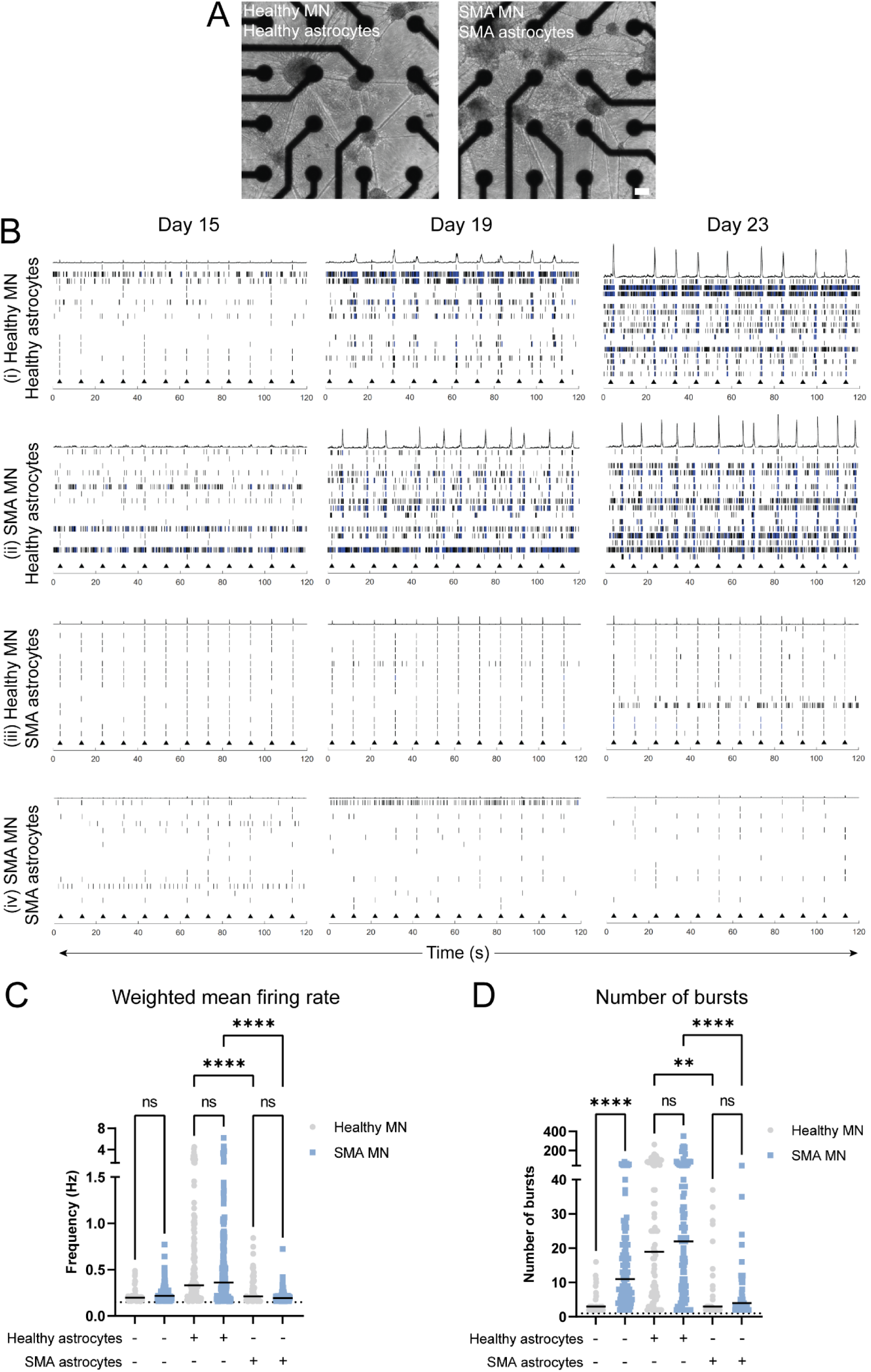
SMA patient iPSC-derived astrocytes diminish evoked motor neuron activity in direct contact co-cultures. **(A)** Brightfield images of motor neuron (MN) and astrocyte co-cultures plated on to 16 electrode CytoView microelectrode (MEA) array plates. Scale bar: 100µm. **(B)** Representative raster plots demonstrating electrically evoked (black triangles) motor neuron activity from (i) healthy MN healthy astrocytes, (ii) SMA MN healthy astrocytes, (iii) healthy MN SMA astrocytes and (iv) SMA MN SMN astrocytes co-cultures at Day 15, 19 and 23. Black lines and blue line clusters represent action potentials/spikes and bursting events on single electrodes, respectively. **(C)** Weighted mean firing rate and **(D)** number of bursts graphs demonstrate combined electrically evoked activity from motor neuron only and motor neuron-astrocyte co-culture recordings (4 minutes) from Day 7-28. Grey circular data points are from healthy motor neurons and blue square data points are from SMA patient-derived motor neurons. Data below the background threshold, determined by low levels of activity from astrocyte only wells, were removed from the analysis (black dashed lines). Statistical testing: Kruskal-Wallis test with Dunn’s multiple comparisons test, ****p-val<0.0001, **pval=0.0013. Data are from 4 independent MEA plates (n=4), ≥3 technical replicates (wells) per condition, 5 independent cell lines (2 healthy and 3 SMA iPSC lines; N=5).

Healthy and patient-derived motor neurons overall showed a similar onset and maturation of evoked spiking and bursting activity in response to stimulation over time (Day 15, 19 & 23) when co-cultured with healthy-derived astrocytes (Figure 2Bi-ii; Figure 2C and 2D, p-val=ns). We did note a significant increase in the number of bursts in the patient-derived motor neuron only cultures when compared to the equivalent motor neuron cultures derived from healthy iPSCs (Figure 2D; ****p-val>0.0001). Interestingly, this increase was not apparent once the SMA motor neurons were co-cultured with the healthy astrocytes (Figure 2D, p-val=ns). In stark contrast, motor neurons co-cultured with patient-derived astrocytes showed minimal activity across all represented time points (Figure 2Biii-iv; Day 15, 19 & 23) and demonstrated statistically significant reductions in weighted mean firing rate (Figure 2C, ****p-val<0.0001) and number of bursts (Figure 2D, **p-val=0.0013, ****p-val<0.0001) when compared to motor neurons cultured with healthy-derived astrocytes. These trends in motor neuron function within the healthy and patient-derived astrocyte co-cultures were also apparent within spontaneous activity recordings (Figure 3A-C). Figure 3A further demonstrates that both healthy and patient-derived motor neurons are capable of generating spontaneous network bursting events (magenta squares) without evoked stimulation when co-cultured with the healthy astrocytes, although higher levels of variability between replicates for each motor neuron-astrocyte condition are observed (Figure 3A; Rep 1 vs Rep 2). Consistent with the evoked datasets, significantly lower levels of spontaneous motor neuron weighted mean firing rate (Figure 3B, ***p-val=0.0005, ****p-val<0.0001) and bursting activity (Figure 3C, *p-val=0.0372, **p-val=0.0012) were observed in the patient astrocyte co-cultures compared to the healthy astrocyte co-cultures.

**Figure 3.**
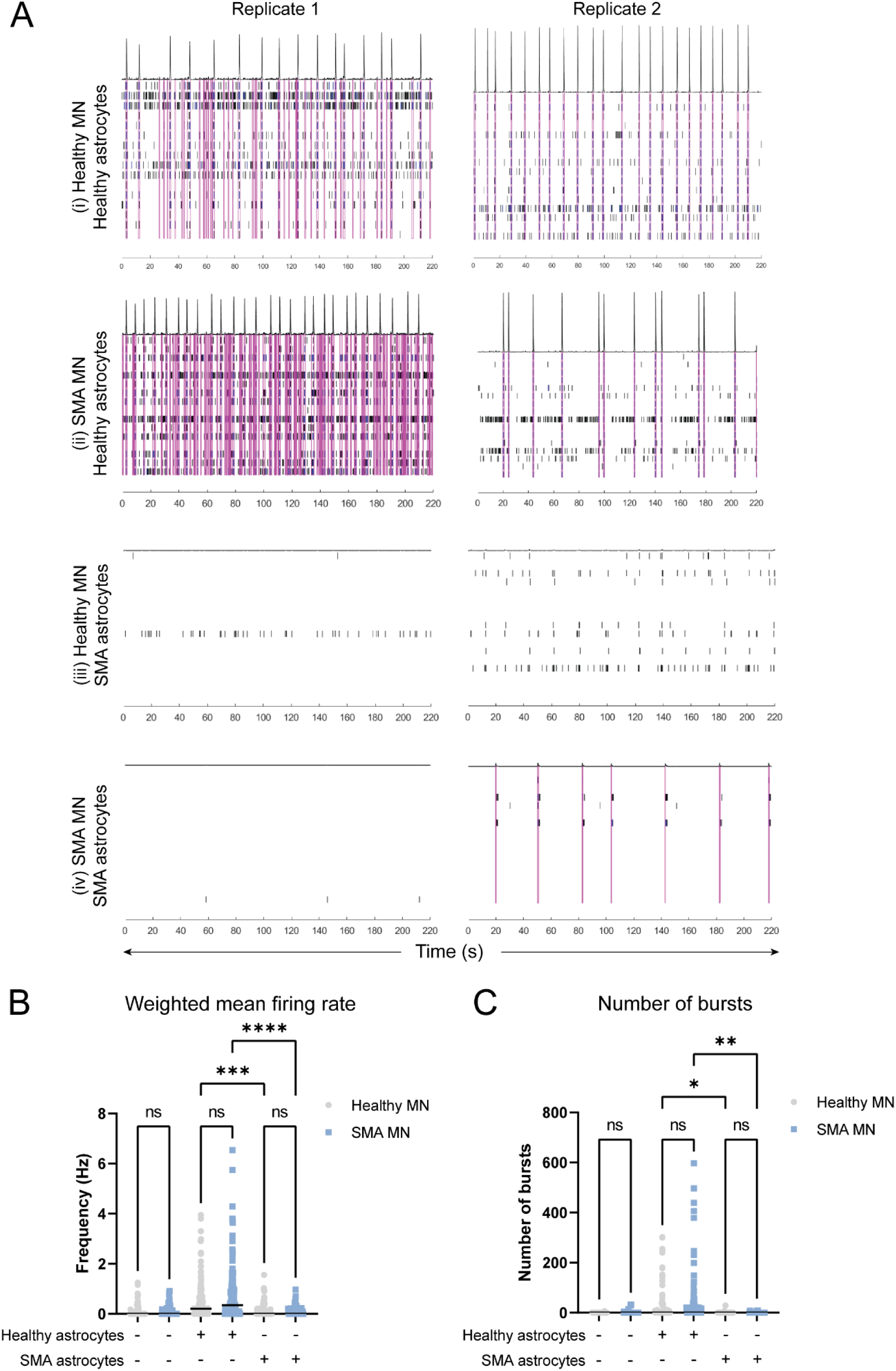
SMA patient iPSC-derived astrocytes diminish spontaneous motor neuron activity in direct contact co-cultures. **(A)** Representative raster plots demonstrating spontaneous motor neuron activity at Day 28 from (i) healthy MN healthy astrocytes, (ii) SMA MN healthy astrocytes, (iii) healthy MN SMA astrocytes and (iv) SMA MN SMN astrocytes co-cultures. Plots from 2 independent replicates are shown per co-culture condition. Black lines and blue line clusters represent action potentials/spikes and bursting events on single electrodes, respectively. Magenta boxes represent organized network bursting events occurring across multiple electrodes without artificial stimulation. **(C)** Weighted mean firing rate and **(D)** number of bursts graphs demonstrate combined spontaneous activity from motor neuron only and motor neuron-astrocyte co-culture recordings (4 minutes) from Day 7-28. Grey circular data points are from healthy motor neurons and blue square data points are from SMA patient-derived motor neurons. Data were excluded from wells that did not produce any motor neuron activity across all recordings (zero data points). Statistical testing: Kruskal-Wallis test with Dunn’s multiple comparisons test, ****p-val<0.0001, ***pval=0.0005, **pval=0.0012, *pval=0.0372. Data are from 4 independent MEA plates (n=4), ≥3 technical replicates (wells) per condition, 5 independent cell lines (2 healthy and 3 SMA iPSC lines; N=5).

### Reduced glutamate transporter (GLAST) levels in SMA patient iPSC-derived spinal cord astrocytes

The transcriptomic and MEA data featured in Figures 1-3 demonstrate that SMA iPSC patient-derived astrocytes modeling cortical or spinal cord regions of the CNS have intrinsic defect in synaptic regulation, with the latter directly able to diminish motor neuron function. *In vivo*, these astrocytes have a fundamental role in glutamate homeostasis at motor neuron afferent synapses via Na^+^ dependent excitatory amino acid transporters, EAAT1/GLAST and EAAT2/GLT-1. Since synaptic and glutamate-related gene ontology terms were enriched within the downregulated transcripts from the RNAseq dataset, we decided to further explore if active glutamate uptake could be defective in SMA astrocytes. Baseline (UTX) and treated (2hr 100µM glutamate treatment) intracellular glutamate levels were measured in astrocyte only cultures. Healthy and SMA patient-derived astrocytes showed increased intracellular glutamate levels after treatment; however, SMA astrocytes had significantly higher glutamate levels at baseline and post-treatment compared to healthy-derived astrocytes (Figure 4A; ****p-val<0.0001). To assess glutamate transporter function, astrocyte cultures were treated with glutamate transporter inhibitor compounds, selective to GLAST (50µM UCPH101) or to GLT-1 (50µM/100µM dihydrokainic acid; DHK) before measuring intracellular glutamate levels. The UCPH101 treatment to inhibit GLAST glutamate uptake led to a significant decrease in intracellular glutamate levels in healthy-derived astrocytes (Figure 4B, ***p-val=0.0003; Figure 4D, 32.6% decrease), which was not observed in SMA patient-derived astrocytes (Figure 4B, p-val=ns; Figure 4D, 7.2% decrease). However, specific inhibition of GLT-1 in both healthy and patient-derived astrocytes showed comparable trends of significantly decreased intracellular glutamate levels at both lower (Figure 4C, **p-val<0.005; Figure 4D, 27.5% and 20.3% decrease) and higher concentrations of DHK treatment (Figure 4D, ****pval<0.0001, Figure 4D, 33.1% and 33.7% decrease). The lack of decreased intracellular glutamate levels in patient-derived astrocytes after specific GLAST inhibition prompted us to check for GLAST protein levels across all astrocyte culture conditions. Immunocytochemistry analysis revealed an extensive population of VIMENTIN+GLAST+ cells across astrocyte cultures derived from both healthy iPSC lines (Figure 4Ei-ii), which was greatly reduced in all astrocyte cultures derived from 3 independent SMA patient iPSC lines (Figure 4Eiii-v; white arrows indicate VIMENTIN+GLAST+ cells).

**Figure 4.**
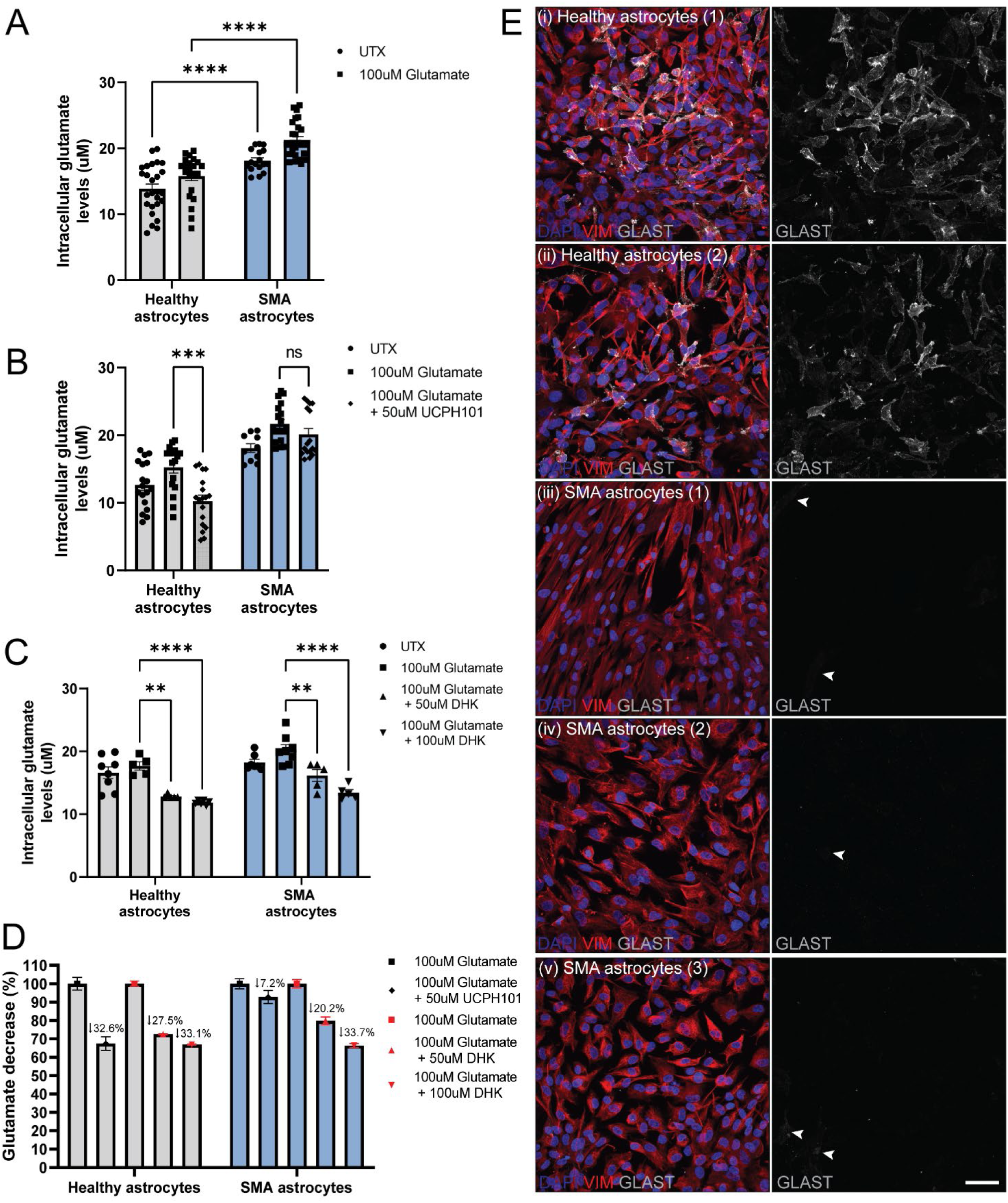
Reduced GLAST glutamate transporter levels in SMA patient iPSC-derived astrocytes. **(A)** Intracellular glutamate levels (µM) at baseline (UTX; black circular data points) and after 100µM glutamate treatment for 2hrs (black square data points) were measured in healthy (grey) and SMA patient (blue) derived astrocytes. In addition to the glutamate treatment, astrocyte cultures were also treated with **(B)** 50µM of the GLAST selective inhibitor, UCPH101 (black diamond data points) or **(C)** 50µM/100µM of the GLT-1 specific inhibitor, dihydrokainic acid (DHK; black triangle data points). **(D)** Intracellular glutamate concentrations after inhibitor treatments were normalized to the levels observed in glutamate treated for healthy and SMA patient-derived astrocyte samples. Relative levels of glutamate decrease are represented as a percentage in the graph. **(E)** Immunocytochemistry analysis of DAPI (nuclei), VIMENTIN (astrocyte intermediate filament) and GLAST (glutamate transporter) was performed on all healthy (i-ii) and SMA patient (iii-v) derived astrocytes. Statistical testing: two-way ANOVA with Tukey’s multiple comparison test, ****pval<0.0001, ***p-val=0.0003, **p-val<0.005. GLAST inhibitor data is from 3 independent experiments (n=3) and GLT-1 inhibitor data is from 2 independent experiments (n=2). 5 independent cell lines (2 healthy and 3 SMA iPSC lines, N=5) were used across all experiments.

### Selective inhibition of GLAST in healthy motor neuron-astrocyte co-cultures recapitulates diminished motor neuron activity observed in SMA patient astrocyte cultures

To assess if the low levels of GLAST protein could contribute to the diminished motor neuron activity observed in patient-derived astrocyte co-cultures, we recorded motor neuron activity in healthy astrocyte co-cultures before and after UCPH101 treatment to mimic the effects of GLAST reduction. Healthy cultures treated with DMSO (vehicle control) showed no changes in motor neuron activity when compared to pre-treated conditions (Figure 5Ai-ii). This is in stark contrast to the diminished motor neuron activity observed after healthy cultures were treated with the GLAST inhibitor, 50µM UCPH101 (Figure 5Bi-ii), which showed comparable low levels of motor neuron activity to those observed within patient-derived astrocyte co-cultures (Figure 5C-D). Specifically, the weighted mean firing rate and number of bursting events of motor neurons from both healthy co-cultures after UCPH101 treatment and patient-derived astrocyte co-cultures were significantly downregulated compared to pre-treated activity (Figure 5E-F, **pval<0.005, ****p-val<0.0001) and showed no statistically significant difference when compared to each other (Figure 5E-F, p-val=ns).

**Figure 5.**
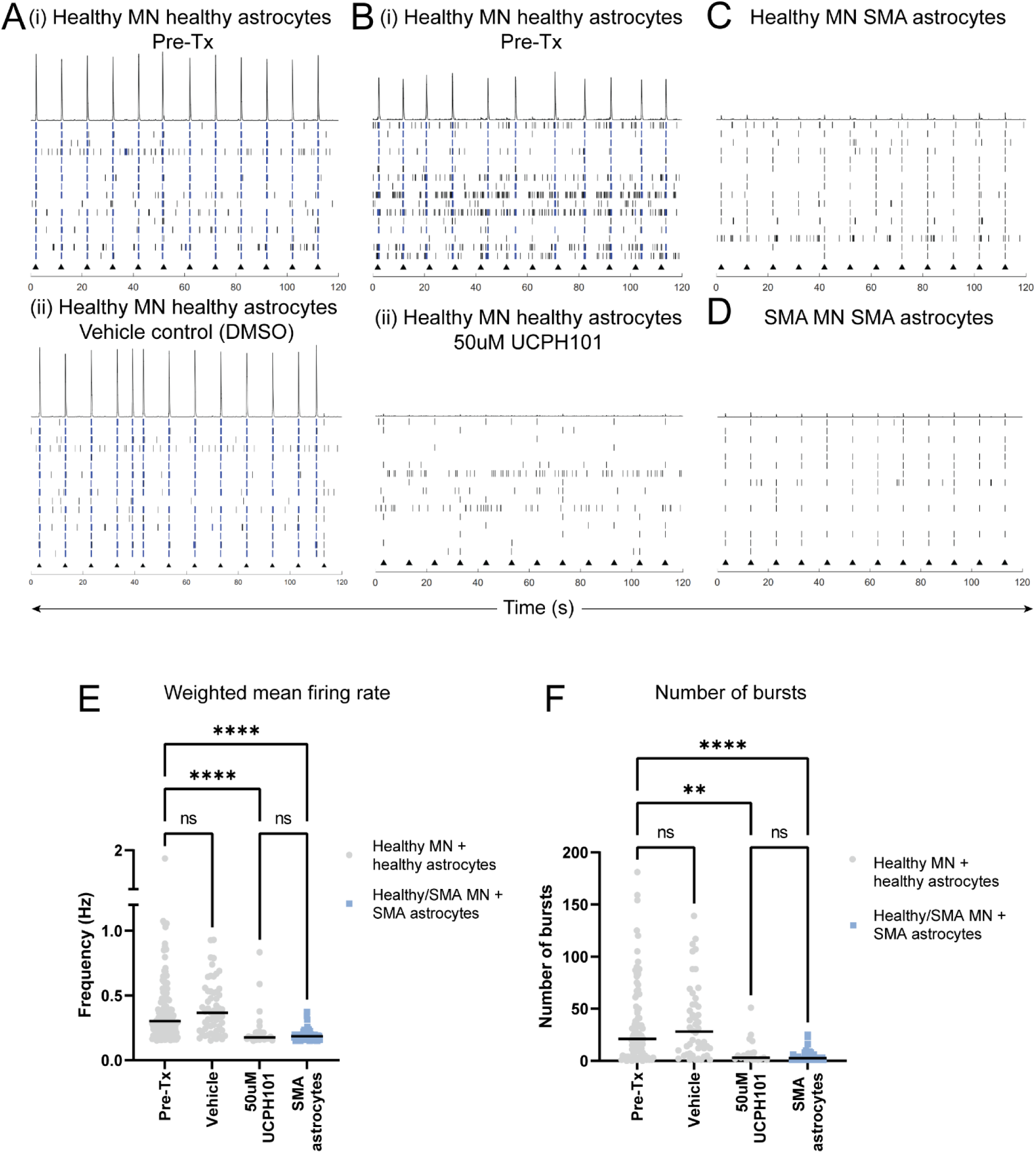
Inhibition of GLAST in healthy cultures mimics diminished motor neuron activity observed in SMA patient-derived astrocyte co-cultures. Raster plots showing evoked motor neuron (MN) activity in **(Ai)** Pre-treatment (Pre-Tx) and **(Aii)** DMSO treated cultures, and **(Bi)** Pre-Tx and **(Bii)** 50µM UCPH101 (selective GLAST inhibitor) treatment, from healthy MN + healthy astrocyte co-cultures at Day 28. Black lines and blue line clusters represent action potentials/spikes and bursting events on single electrodes, respectively. Raster plots in **(C)** and **(D)** show motor neuron activity in healthy MN + SMA astrocyte and SMA MN + SMA astrocyte co-culture, respectively. Motor neuron **(E)** weighted mean firing rate and **(F)** number of bursts data are show from all co-culture conditions (grey circular data are from healthy MN + healthy astrocytes, blue square data are from healthy/SMA MN + SMA astrocyte co-culture) recorded from Day 13-19. Statistical testing: Kruskal-Wallis test with Dunn’s multiple comparisons test, ****p-val<0.0001, **pval<0.005. Data are from 2 independent MEA plates (n=2), ≥3 technical replicates (wells) per condition, 5 independent cell lines (2 healthy and 3 SMA iPSC lines; N=5).

### Increased SMN protein levels in SMA patient iPSC-derived astrocytes partially restores GLAST expression and function

The downregulation of genes associated with astrocyte-mediated glutamate homeostasis within patient-derived samples suggests that the GLAST phenotype is likely to be SMN-dependent. To test this hypothesis, we assessed the levels of GLAST protein restoration in *SMN1* corrected patient astrocytes. Lentiviral DNA constructs containing FLAG tagged human *SMN1* (SMN:FLAG) or a GFP reporter downstream of human *SMN1* (SMN:GFP) were used in this study. As expected, immunocytochemistry analysis revealed the reduction of SMN protein in patient-derived astrocytes when compared to healthy astrocytes (Figure 6A-B), and SMN protein was readily detected in patient astrocytes transduced with the SMN:FLAG (Figure 6C) and SMN:GFP (Figure 6D) lentiviruses.

**Figure 6.**
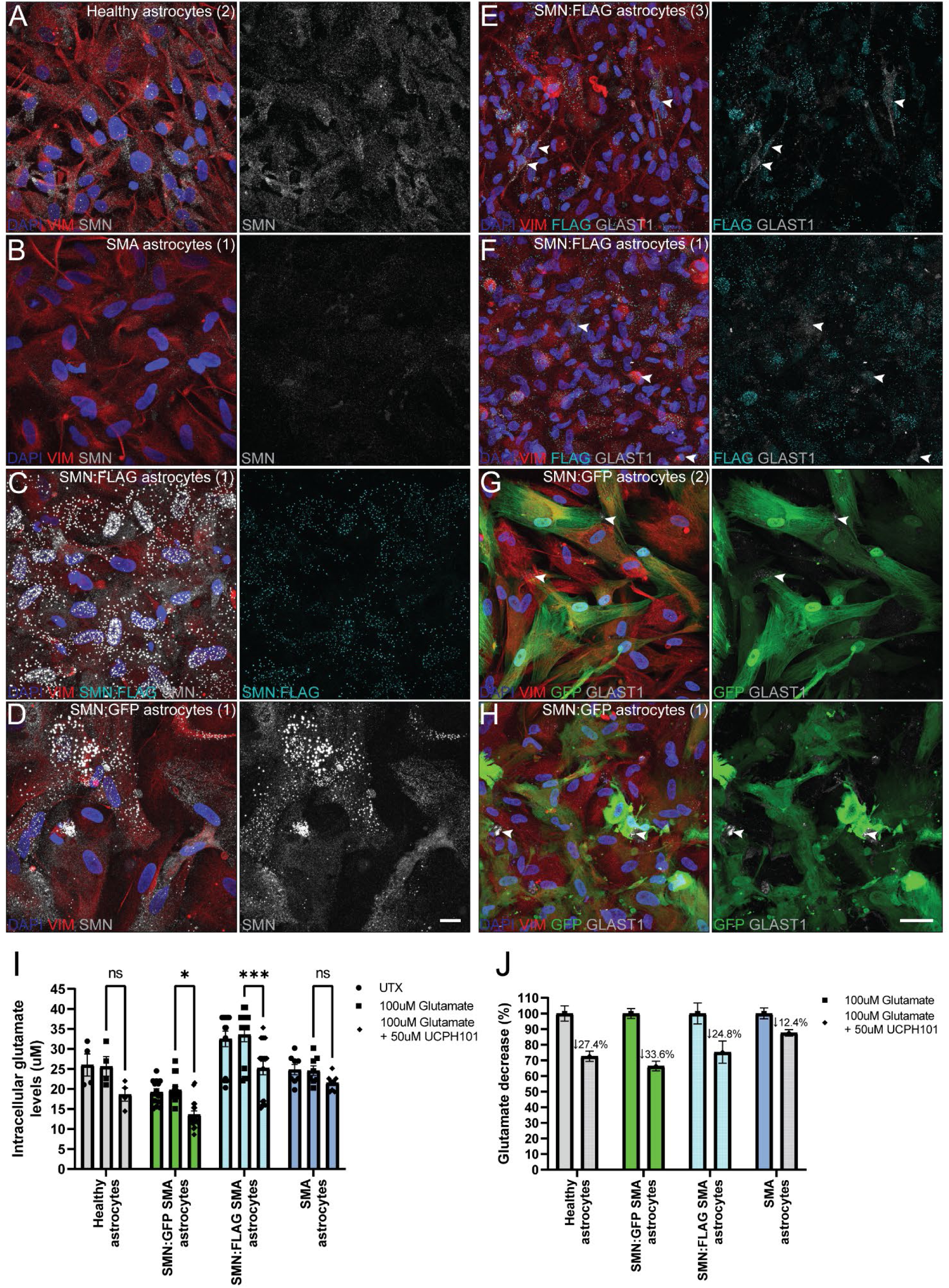
Lentiviral mediated delivery of SMN into SMA patient-derived astrocytes leads to a partial restoration of GLAST protein levels. Immunocytochemistry analysis performed on **(A)** healthy astrocytes, **(B)** SMA astrocytes, **(C)** SMN:FLAG and **(D)** SMN:GFP transduced SMA astrocytes assessing DAPI (nuclei), VIMENTIN (astrocyte intermediate filament) and SMN protein levels. Signal from anti-FLAG antibody is shown in **(C)**. Endogenous GFP signal from SMN:GFP transduced SMA astrocytes **(D)** could not be detected during SMN immunocytochemistry analysis due to the acetone/methanol fixation method required to detect SMN protein. A similar analysis was completed on **(E&F)** SMN:FLAG and **(G&H)** SMN:GFP transduced SMA astrocytes to assess GLAST protein. PFA fixation allowed for the detection of FLAG and GFP signal for this analysis **(E-H)**. White arrows point to GLAST+ VIMENTIN+ FLAG+/GFP+ cells in SMN transduced SMA astrocyte images. Scale bar for A-D: 20µm, scale bar for E-H: 50µm. **(I)** Intracellular glutamate levels were assessed across healthy (grey bars), SMN:GFP SMA astrocytes (green bars), SMN:FLAG SMA astrocytes (cyan bars) and SMA patient-derived astrocytes (blue bars). Data is shown from untreated (UTX; black circular data points), 100µM glutamate treated (black square data) and 100µM glutamate + 50µM UCPH101 treated (clack diamond data points) for each astrocyte condition. **(J)** Relative levels of glutamate decrease (normalized to 100µM glutamate for each astrocyte dataset) are represented as a percentage in the graph. Statistical testing: two-way ANOVA with Tukey’s multiple comparison test, ***p-val=0.0005, *p-val=0.04. SMN:FLAG and SMN:GFP SMA astrocyte data were collected from 3 independent SMA astrocyte lines, 2 differentiation replicates per line (N=6). All other data were collected from 5 independent cell lines (2 healthy and 3 SMA iPSC lines, N=5, 2 differentiation replicates per line).

Immunocytochemistry analysis of GLAST protein in the SMN-restored patient astrocytes showed differing levels of GLAST restoration and localization within the cells (Figure 6E-H). GLAST+ puncta co-localizing with SMN:FLAG+ (Figure 6E) or SMN:GFP+ (Figure 6G-H) cells were observed in some of the SMN-restored patient astrocyte cultures, showing a similar staining pattern to that observed in healthy-derived cultures. Others showed a more diffuse, cytoplasmic GLAST+ signal in SMN-restored patient astrocytes (Figure 6F). To assess if this increase in GLAST protein levels via SMN re-expression could also improve GLAST-mediated glutamate functioning, intracellular glutamate levels were assessed in SMN-restored patient astrocytes before and after 50µM UCPH101 inhibitor treatments. A significant decrease in intracellular glutamate levels was observed across all SMN-restored patient astrocyte lines when treated with UCPH101 (Figure 6I, *p-val=0.04, ***p-val=0.0005), which showed a similar percentage of intracellular glutamate decrease when compared to the UCPH101 treated healthy-derived astrocyte samples (Figure 6J, healthy: 27.4% decrease, SMN:GFP: 33.6% decrease, SMN:FLAG: 24.8% decrease, SMA astrocyte: 12.4% decrease).

### Abnormal increase in caveolin-1 protein levels in SMA patient iPSC-derived astrocytes is partially rescued by SMN lentiviral transduction

To further unravel the disease mechanism involving SMN and GLAST deficiency in patient astrocytes, we assessed protein levels of caveolin-1 (CAV-1), an SMN-interacting lipid raft scaffold protein involved in caveolae-mediated cell signaling, endocytosis and glutamate transporter regulation in astrocytes (49). Using immunocytochemistry analysis, we found an increase in CAV-1 protein levels and large CAV-1+ accumulations in SMA patient-derived astrocytes compared to healthy controls (Figure 7A-B). Interestingly, this CAV-1 phenotype was less apparent in patient astrocytes transduced with SMN:FLAG (Figure 7C) or SMN:GFP (Figure 7D). These trends in CAV-1 levels could also be corroborated via immunoblot analysis (Figure 7E-G). Patient-derived astrocytes had consistently higher levels of CAV-1 (****p-val<0.0001; 207% increase), an increase which was also observed in patient lumbar spinal cord samples (58% increase), while SMN re-expression in patient-derived astrocytes caused a significant decrease in CAV-1 levels (SMN:GFP: ****p-val<0.0001, 162% decrease; SMN:FLAG: **p-val=0.0039, 110% decrease) (Figure 7F-G). Interestingly, we noted an inverse correlation specifically between SMN:GFP and CAV-1 levels; individual astrocyte replicates with higher levels of SMN:GFP signal showed lower levels of CAV-1 signal (Figure 7H). This prompted us to assess the lentivirus transduction efficiency via immunocytochemistry analysis by calculating the percentage of DAPI+ VIMENTIN+ GFP/FLAG+ cells within the total cell population counted (Figure 7I), which demonstrated a varying levels of lentivirus transduction rates across replicate lines (approximately 20-60%). However, these data highly aligned with immunoblot GFP/CAV-1 signal observation, validating that astrocyte lines with higher SMN lentivirus transduction efficiency had lower levels of CAV-1.

**Figure 7.**
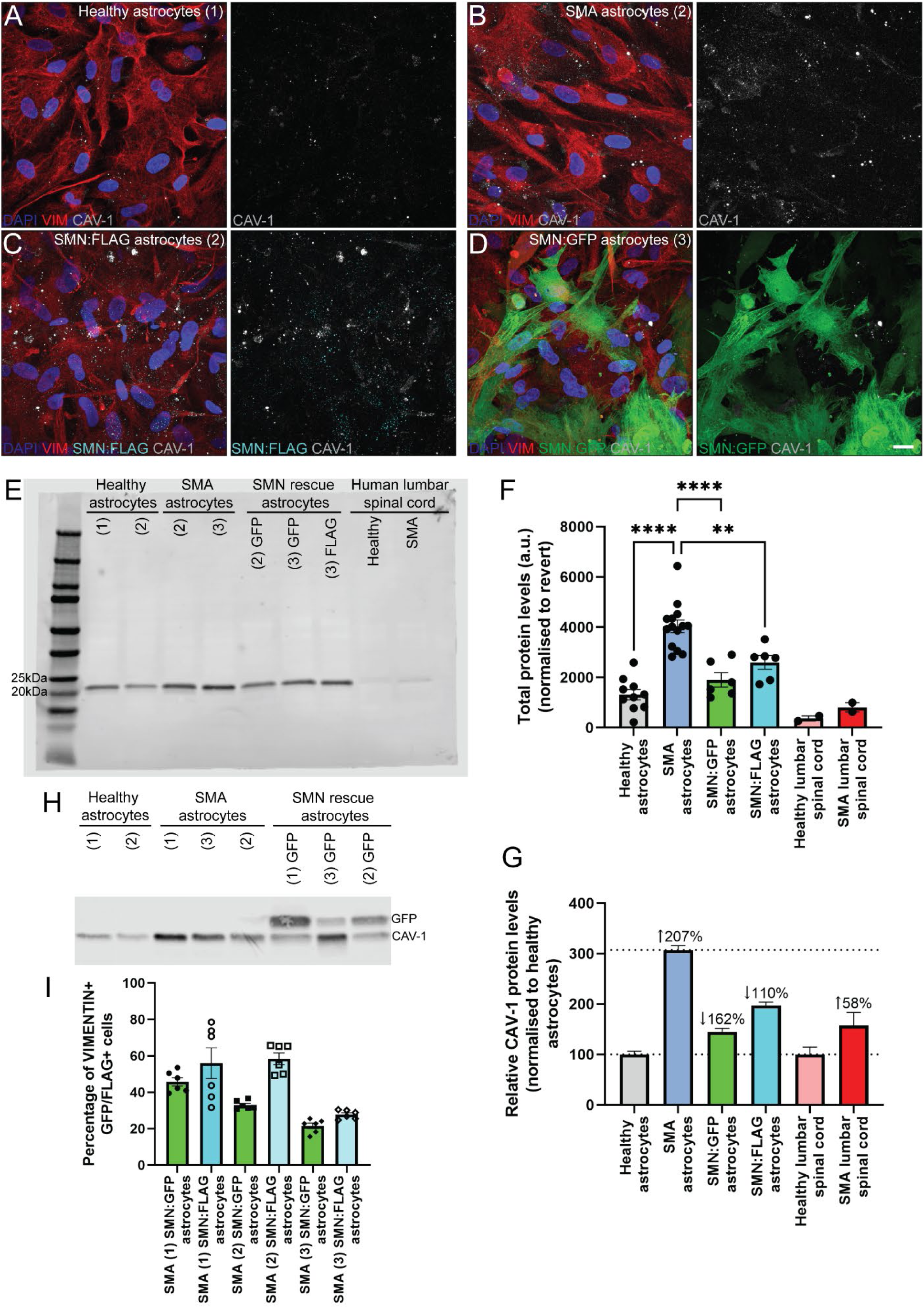
Increased caveolin-1 protein levels in SMA patient iPSC-derived astrocytes can be partially corrected via SMN restoration. Immunocytochemistry analysis of DAPI (nuclei), VIMENTIN (astrocyte intermediate filament) and caveolin-1 (CAV-1) protein levels in **(A)** healthy astrocytes, **(B)** SMA astrocytes and SMA astrocytes transduced with **(C)** SMN:FLAG or **(D)** SMN:GFP lentiviruses. **(E)** CAV-1 immunoblotting was performed on healthy, SMA patient and SMA patient SMN:GFP/FLAG transduced astrocytes (iPSC-derived samples), in addition to post-mortem lumbar spinal cord tissues from healthy individuals and SMA patients. **(F)** Total (measured in arbitrary units (a.u.) and **(G)** relative CAV-1 protein levels were quantified across technical replicates. SMA and SMA SMN transduced iPSC-derived samples were normalized to healthy iPSC-derived astrocytes; SMA lumbar spinal cord protein levels were normalized to healthy lumbar spinal cord samples. **(H)** CAV-1 and GFP protein levels were assessed in healthy, SMA patient, and SMA patient SMN:GFP transduced astrocyte samples via immunoblotting. **(I)** Lentiviral transduction efficiency was assessed by calculating the percentage of VIMENTIN+ (astrocyte intermediate filament) and GFP/FLAG+ cells out of the total cell population (VIMENTIN+ DAPI+). Statistical testing: one-way ANOVA with Tukey’s multiple comparison test, ****p-val<0.0001, **p-val=0.0039. SMN:FLAG and SMN:GFP SMA astrocyte data were collected from 3 independent SMA astrocyte lines, 2 differentiation replicates per line (N=6). All other data were collected from 5 independent cell lines (2 healthy and 3 SMA iPSC lines, N=5, 4-5 differentiation replicates per line).

### Minor improvements in motor neuron spike amplitude observed in SMN transduced patient-derived astrocyte co-cultures

Since we observed improvements in the GLAST and CAV-1 phenotypes after SMN restoration in patient-derived astrocytes, we co-cultured these astrocytes with healthy-derived motor neurons to assess if the diminished motor neuron activity phenotype could also be rescued. Figure 8A-C show representative brightfield images demonstrating equivalent electrode coverage of motor neurons across the different co-culture conditions. As expected, we were able to readily detect spikes and bursting activity in response to electrical stimulation in purely healthy co-culture conditions (Figure 8Di), which was severely diminished in SMA patient-derived co-cultures (Figure 8Dii). In SMN:GFP and SMN:FLAG astrocyte co-cultures, motor neuron activity was still minimal with no improvements observed in spike frequency nor bursting activity (Figure 8Dii-iv). This same observation was noted across spontaneous and electrically evoked motor neuron activity in which SMN restored co-cultures showed comparable weighted mean firing rates (i) and number of bursts (ii) to patient-derived co-cultures (Figure 8E-F, p-val=ns). We did note a slight increase in the spike amplitude upon motor neuron stimulation in the SMN transduced patient-derived astrocyte co-cultures (Figure 8Dii-iv; red arrows) compared to patient-derived cultures (Figure 8Dii; red arrow). However, the spike amplitude did not reach the same levels as those observed in the healthy co-cultures (Figure 8Di).

**Figure 8.**
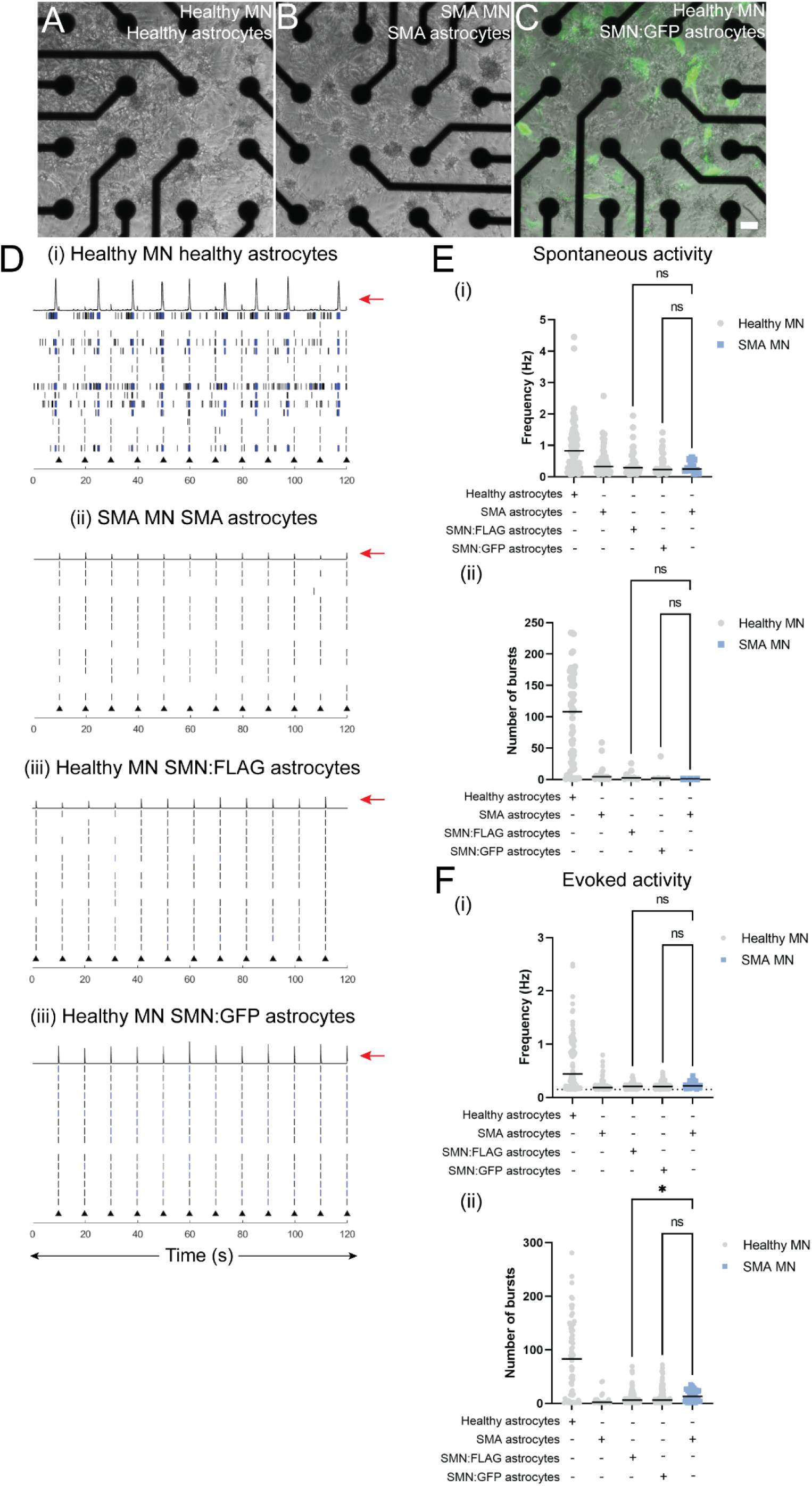
Improvements in spike amplitude, but minimal activity observed in motor neurons co-cultured with SMN transduced SMA patient derived astrocytes. Brightfield images across healthy **(A)**, SMA **(B)** and SMN re-expressing **(C)** motor neuron astrocyte co-cultures on MEA plates. Scale bar: 100µm. **(Di-iv)** Raster plots demonstrating electrically evoked (black triangles) motor neuron activity at Day 17 of co-culture across the different astrocyte conditions. Black lines and blue line clusters represent action potentials/spikes and bursting events on single electrodes, respectively. Red arrows highlight differences in spike amplitude. Spontaneous **(Ei-ii)** and electrically evoked **(Fi-ii)** motor neuron activity (weighted mean firing rate and number of bursts) across different astrocyte co-culture conditions. Grey circular data points are from healthy motor neurons and blue square data points are from SMA patient-derived motor neurons. Statistical testing: Kruskal-Wallis test with Dunn’s multiple comparisons test, p-val=ns. Motor neuron data were collected from astrocyte co-cultures derived from 2 healthy iPSCs, 3 SMA iPSC lines, 12 SMN transduced patient astrocyte differentiations. Data were collected from 2 MEA plates total.

## Discussion

Using a clinically relevant model system, the main aims of our study were to define i) molecular candidates with a striking phenotype in SMA patient-derived astrocytes, ii) assess phenotype association with SMN deficiency and iii) determine the contribution of the astrocytic-intrinsic phenotype to motor neuron activity. In this study, we highlight the overlooked role of glial-intrinsic defects that can mediate human motor neuron dysfunction in SMA specifically through impaired glutamate neurotransmission regulation. While SMN delivery via lentivirus transduction improved the GLAST and CAV-1 phenotype in SMA astrocytes, it was not sufficient to restore motor neuron activity in patient-derived astrocyte cultures. This may indicate that greater modulation of SMN levels is needed to reverse astrocytic disease phenotypes or that other astrocytic defects, such as reactivity instigating irreversible cellular defects, may be insensitive to SMN restoration. Either outcome will have important implications for current SMN therapies.

### Assessment of molecular candidates driving intrinsic defects in SMA astrocytes: GLAST

Transcriptome analysis of human SMA patient-derived astrocytes revealed for the first time a striking downregulation of molecular candidates associated with astrocytic-mediated modulation of the synapse. In addition to corroborating other studies which previously found decreased expression of astrocyte-specific ion channels (K_ir_4.1), glutamate transporter (EAAT1) and ephrins (34, 35), our data highlight additional molecular candidates which may further disrupt human astrocytic regulation of the synapse and neural synaptogenesis. Our study also suggests that human SMA astrocytes are likely to exacerbate the disrupted glutamate neurotransmission phenotype previously associated with proprioceptive innervation of motor neurons (10). GLAST and GLT-1 are thought to be the two specific astrocyte transporters responsible for glutamate uptake and clearance from the synaptic cleft (54). Specifically, we validated the decreased levels of GLAST within forebrain and spinal cord patterned astrocytes derived from SMA patient Type I and II patient iPSC lines, suggesting that this a consistent phenotype across multiple regions of the CNS and across different SMA disease severities. The GLT-1 phenotype was inconsistent across astrocyte populations, with decreased GLT-1 expression observed within forebrain-like astrocytes, but preserved GLT-1-function in spinal cord-like astrocytes. This observation is not too surprising since astrocyte glutamate transporter expression differs across the CNS (55-57). Despite the decreased levels of GLAST, it was surprising to observe increased intracellular glutamate levels in patient-derived astrocytes, which contrasts with the decreased glutamate uptake observations from SMN siRNA mouse-derived astrocyte cultures (34).

We hypothesize that species differences and genetic derivation (SMN siRNA vs *SMN1* mutation) of the cells may explain this discrepancy. It is also highly possible that GLAST1 defects are coupled with disrupted glutamate-glutamine metabolism or plasma membrane integrity which also affect astrocytic intracellular glutamate levels, as previously observed in Alzheimer’s disease (AD) (58, 59) and Parkinson’s disease (49). In relation to other motor neuron diseases, we find that our study contrasts to observations in amyotrophic lateral sclerosis patients and animal models (60-62), and animal models of spinal cord motor neuron degeneration (63, 64) in which low levels and impaired function of GLT-1 specifically drives impaired glutamate clearance. Similar to our study, GLAST-specific impairment has been reported for other neurological diseases, including ADHD (65) and episodic ataxia (66). In broader CNS disorders, GLAST and GLT-1 dysfunction is typically observed, such as in AD pathology (67), but contradictory evidence regarding increased or decreased levels of glutamate transporters exists for other disorders including Parkinson’s disease (49, 68) and multiple sclerosis (69, 70). This variability indicates the specific genetic aberration underlying neurological diseases may impact glutamate transporter expression differently. We therefore hypothesize that the regulation and expression of GLAST may be more sensitive to SMN deficiency than GLT-1 within SMA spinal cord astrocytes.

### The link between the GLAST phenotype and SMN deficiency: caveolin-1

In addition to the GLAST phenotype, caveolin-1 (CAV-1), an SMN-interacting protein found within plasma membrane lipid rafts (46, 47) with a previously reported role in astrocytic glutamate transporter regulation (49), showed a striking upregulated phenotype in SMA patient-derived astrocytes. This is in contrast to the decreased CAV-1 levels previously reported in SMA Type I patient fibroblasts (46). It is worth noting caveolin proteins are heavily understudied in the CNS, and caveolae distribution and mediated endocytosis varies greatly across different tissues and cell types (71). It is also likely that CAV-1 regulation of cellular cholesterol levels and distribution (72, 73) to facilitate endocytic pathways (71) may have a greater role within astrocytic cholesterol biosynthesis and metabolism compared to fibroblasts. Increasing SMN levels improved the severely reduced GLAST and increased CAV-1 phenotypes in SMA patient-derived astrocytes. This overall suggests a mechanism involving SMN-CAV-1 regulation of GLAST expression in astrocytes, which draws similarities with a previous study showing DJ-1 deficiency in Parkinson’s disease astrocytes causes defective lipid raft-dependent endocytosis (via flotillin and caveolin-1) and glutamate transporter expression (GLT-1) (49). However, SMN re-expression only led to a partial restoration of the GLAST and caveolin-1 phenotypes. We noted some variability in the SMN lentivirus transduction efficiency (20-60%) across the different patient iPSC lines which may explain the partial phenotype rescue and minimal functionality in motor neuron co-cultures. Successfully transduced astrocytes however did show intense SMN+ signal, even more so than the healthy-derived astrocytes. Further work is needed to address if specific modulation of SMN restoration (across total population and levels in individual cells) is needed to counteract the disease mechanism or whether other astrocyte intrinsic changes, such as irreversible A1-reactivity (74), could be insensitive to SMN restoration. In addition to CAV-1, our RNA seq data indicate mis-expression of metabotropic glutamate receptors and Na^+^/K^+^ ATPases within patient-derived astrocytes, which may additionally contribute to defective glutamate transporter expression and function (75-77). It is possible that abnormal Notch signaling (78) and calcium signaling (31) previously observed in SMA astrocytes, or even neural-driven changes in neurotransmission (79, 80) may exacerbate glutamate transporter defects in SMA. Indeed, alteration of CAV-1 and the lipid raft-scaffold supporting local protein translation and endocytosis of cell surface proteins, such as GLAST, in SMN deficiency may also cause problems for astrocyte-mediated neuromodulation involving rapid protein surface diffusion and plasma membrane protein turnover in response to neural activity (81).

### The impact of astrocytic-intrinsic defects on motor neuron function in SMA

Our co-culture enabled the separate derivation of astrocytes and motor neurons from healthy and SMA patient iPSC lines, which allowed us to interrogate motor neuron-intrinsic and astrocytic-mediated dysfunction. A key finding of this study was the severely diminished motor neuron activity specifically in patient-derived astrocyte direct contact co-cultures, a phenotype consistent with a previous study using SMA mouse-derived cultures (35). Patient-derived motor neurons were capable of high levels of activity in healthy-derived astrocyte co-cultures. Without the addition of astrocytes, patient-derived motor neurons showed slightly higher firing rate levels and significantly more bursting events than healthy-derived motor neurons, which recapitulates the increased membrane excitability phenotype observed in previous studies (10, 16, 19-21). Strikingly, this phenotype appears to be normalized through the addition of healthy-derived astrocytes. These data together provide unprecedented evidence that astrocytes can drive motor neuron dysfunction in SMA and highlight the importance of restoring astrocyte-mediated neuromodulation during disease.

Importantly, we show that selective GLAST inhibition in healthy-derived co-cultures via the addition of UCPH101, which acts to block GLAST function and glutamate uptake (82), is enough to phenocopy diminished motor neuron activity mediated by SMA patient-derived astrocytes. Similar decreased motor neuron activity has been described within SMA animal models proposing proprioceptive-derived defects impede glutamate neurotransmission (24) and through the addition of generic glutamate transporter inhibitor, TBOA, within iPSC-derived spinal cord-derived co-cultures (83). Deficits in glutamate neurotransmission and reduced levels of GLAST have also been previously associated in AD pathology (84). The molecular consequences leading to reduced motor neuron activity that is instigated by astrocytic glutamate transporter depletion remains to be resolved. It could be hypothesized that reduced astrocytic GLAST would prevent glutamate uptake causing glutamate excitotoxicity at the synaptic cleft and eventually leading to neuronal cell death as described in other neurological diseases (85-87). It is also possible that dysfunctional glutamate uptake within astrocytes could in turn impact motor neuron receptor and transporter expression to decrease overall excitatory neurotransmission, similar to the proprioceptive neuron-derived mechanism in SMA deficiency, which leads to the loss of Kv2.1 ion channel expression in motor neurons (24).

Overall, our work defines a novel SMN-associated disease mechanism involving abnormal glutamate transporter activity and regulation in astrocytes that drives motor neuron inactivity in SMA. *In vivo*, astrocytic-driven dysfunction is likely to occur in combination with intrinsic motor neuron defects and sensory neuron-mediated dysfunction, with potential input from other glial cell types including microglia (15, 50, 88). Studies have already begun to test the efficacy of therapeutic compounds in order to increase motor neuron activity at the central afferent synapse (89, 90). As novel therapies for SMA progress and long-term effects of FDA-approved treatments are assessed in patients, our work highlights the importance of tracking improvements in glial health and glial-mediated neuromodulation, which may be critical to fully restore sensory-motor circuit functionality.

## Acknowledgements

We would like to thank Benjamin O’Brien and the Rao laboratory at the Medical College of Wisconsin (MCW)/Versiti Blood Research Institute (BRI) for providing assistance with the mRNA sequencing experiments and access to the Illumina NextSeq Sequencer. Lentiviral DNA constructs used in this study were kindly provided by the laboratories of Dr. Xue Jun Li (University of Illinois-Chicago) and Dr. Christian Lorson (University of Missouri). We also acknowledge the Viral Vector Core Facility at the MCW/BRI for generating the lentiviruses used in this study and the Cardiovascular Center equipment core at MCW for use of the MEA system. Human spinal cord tissue was obtained from the NIH NeuroBioBank. This work was supported by the Audrey Lewis Young Investigator Award funded by Cure SMA (E.W.) and R21NS102911 (A.D.E)

## Author contribution

E.W. Conception and design, collection and assembly of data, data analysis and interpretation, manuscript writing, financial support. A.D.E. Conception and design, data analysis and interpretation, final manuscript editing, financial support.

